# Ordered and disordered regions of the Origin Recognition Complex (ORC) combine to direct in-vivo binding at symmetric and non-symmetric motif sequences

**DOI:** 10.1101/2024.01.27.577596

**Authors:** Michal Chappleboim, Segev Naveh-Tassa, Miri Carmi, Yaakov Levy, Naama Barkai

## Abstract

The Origin Recognition Complex (ORC) seeds replication-fork formation by binding to DNA replication origins, which in budding yeast contain a 17bp DNA motif. High resolution structure of the ORC-DNA complex revealed two base-interacting elements: a disordered basic patch (Orc1-BP4) and an insertion helix (Orc4-IH). To define the ORC elements guiding its DNA binding *in-vivo*, we mapped genomic locations of 38 designed ORC mutants, revealing that different ORC elements guide binding at different sites. At silencing-associated sites lacking the motif, ORC binding and activity were fully explained by a BAH domain. Within replication origins, we reveal two dominating motif variants showing differential binding modes and symmetry: an asymmetric motif whose binding requires Orc1-BP4 and Orc4-IH, and a symmetric one where another basic patch, Orc1-BP3, can replace Orc4-IH. Disordered basic patches are therefore key for ORC-motif binding *in-vivo*, and we discuss how these conserved, minor-groove interacting elements can guide specific ORC-DNA recognition.

## Introduction

In eukaryotes, DNA replication is initiated by firing hundreds of replication origins across the genome^1–6^. The Origin Recognition Complex (ORC) binds replication origins and recruits the replication machinery. In budding yeast, ORC recognizes a consensus 17-bp T-rich motif^7–16^ while in higher eukaryotes, ORC binding sites lack an apparent motif but are still enriched with T-stretches^17–21^.

A high-resolution cryo-EM structure of the *S. cerevisiae* ORC bound to DNA is available^22^, showing that all six ORC subunits are engaged in backbone interactions. Only two elements, however, formed specific base contacts: an insertion helix within Orc4 (Orc4-IH) and a basic patch (Orc1-BP4) within Orc1. These ORC-DNA specific contacts, in turn, were limited to five of the seventeen base-pair motif sequence^22^. Orc4-IH is unique to budding yeast and its close phylogenetic relatives^23,24^, whereas basic patches, defined as >5 basic residues within a 10-14 AA stretch, are conserved features of Orc1 N-terminus across eukaryotes^22^, although with varying sequences and locations.

Inferring principles of *in-vivo* binding from structural data is challenging. First, ORC binds to hundreds of genomic sites with different motif variants, while the structure describes only one bound sequence. Second, *in-vitro* analysis often relies on truncated proteins that are easier to express and purify. In the ORC-DNA structure, for example, Orc1 lacks most of its disordered N-terminus (Fig. 1A)^22^. Third, chromatin and other accessory proteins that could influence *in-vivo* binding are missing from the structure. Finally, cryo-EM structures capture only conformations of sufficient stability, while genomic preferences could depend on contacts that are required transiently for reaching the stable configuration.

**Figure 1.**
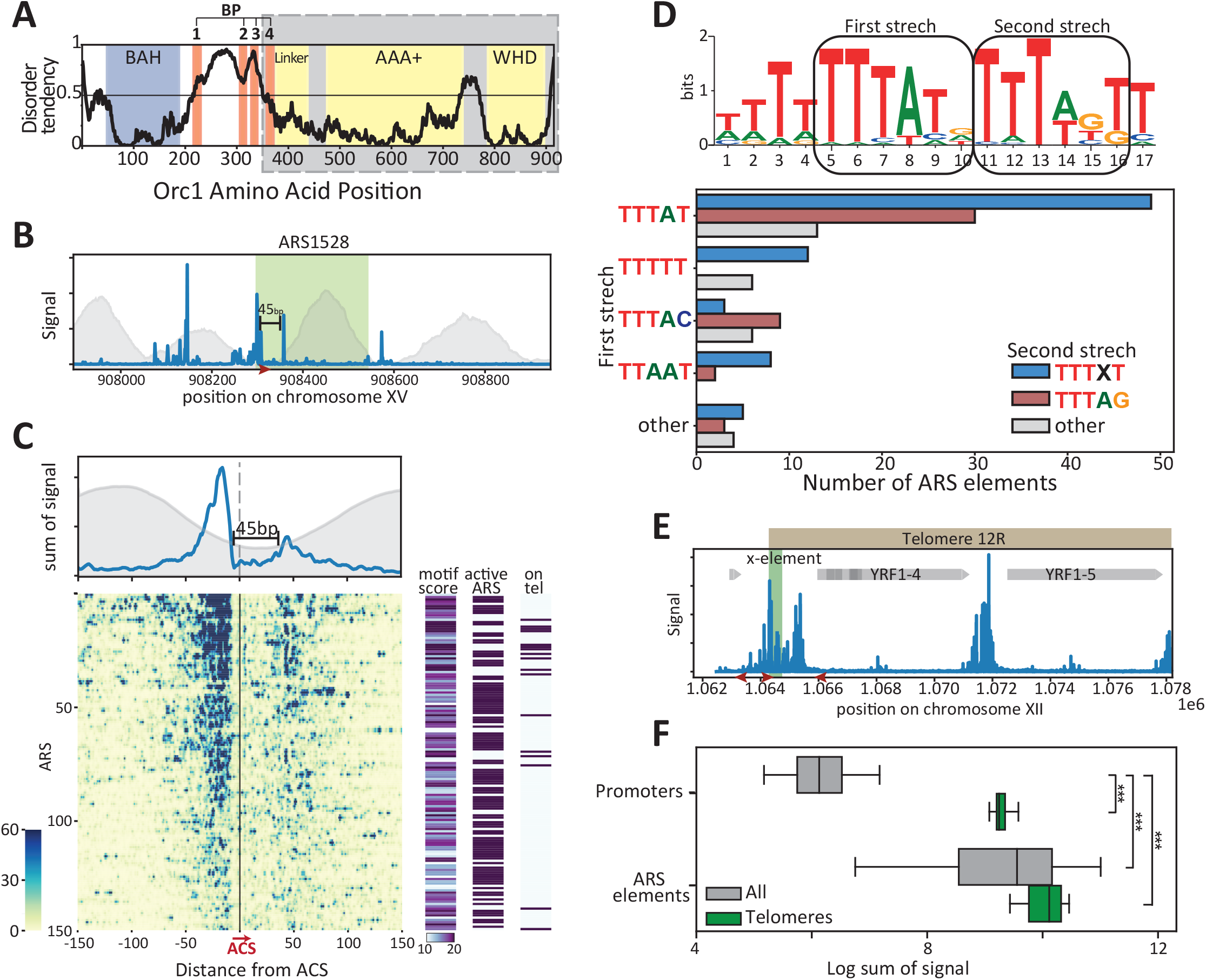
Genome-wide mapping of ORC binding locations reveals two prevailing motif sequences that dominate top-bound origins. *A. Predicted disorder tendency of S*.*cerevisiae Orc1*: The black line indicates the predicted disorder tendency along the Orc1 protein sequence, as calculated by IUPred ^33,34^. Values above 0.5 are considered disordered. Colored regions indicate characterized domains (BAH: bromo-adjacent homology, BP: 4 basic patches, AAA+: ATPase associated activity, WHD: winged helix domain). The cryo-EM structure^22^ was solved using a truncated Orc1, containing the 355-914 AA (Gray). *B. Orc1 binding at a representative origin* (ARS1528): Shown is the normalized binding signal of Orc1 mapped to the indicated genomic segment containing a defined origin (see Methods for read alignment). The red arrow indicates the position and orientation of the ORC consensus sequence. Nucleosome occupancy is indicated in gray (data from Chapal et al.^35^). *C. Orc1 binding across replication origins (ARSs):* Origins and their surrounding 300bp segments were aligned and oriented by the motif sequence. Shown are the average signal (Top) and the mapped reads (Bottom) of the 150 top-bound origins, ordered by the total Orc1 reads. The right columns depict the motif score and ARS replication activity (by Yabuki et al.^36^) of each sequence, and whether it localizes to a telomere. *D. Top-bound origins contain two dominating motif variants:* the ORC motif by YeTFaSCo^37^(Top) and the count of different motif variants present (Bottom) among the 150 top-bound origins. *E. Orc1 binding at the 12R telomere:* Shown is the Orc1 binding signal along the indicated region. The x-element is indicated by a green area. Red arrows indicate the position and orientation of ORC binding motifs, and gray arrows indicate open reading frames (ORFs). Note the strong binding signal at the motif bordering the x-element, and at the promoters of the silenced YRF1-4 and YRF1-5 genes. *F. Orc1 binds gene promoters on telomeres:* Shown are the distributions of Orc1 binding at origins and promoters found away or near telomeres, as indicated. (Distributions shown as box-plots, see methods).

Here, we defined ORC elements required for its *in-vivo* DNA binding by mapping the genomic binding of 38 designed ORC mutants. We describe three classes of ORC binding sites that depend on distinct ORC elements. The first class comprises motif-lacking sites, where ORC is involved in gene silencing^25^, and this binding is fully explained by Orc1 bromo-adjacent homology (BAH) domain. The other two classes include strongly bound replication origins, which we find to sub-divide based on two dominating motif variants: a symmetric motif containing a proximal TTTAT stretch and a distal TTTxT stretch, and an asymmetric motif where the distal stretch is replaced by TTTAG. Orc1-BP4 is necessary for binding at both motifs, while Orc4-IH is required for the asymmetric one, but can be replaced by another basic patch, Orc1-BP3, at the symmetric variant. We discuss the possible implications of our results for the mechanism and ORC-DNA binding *in-vivo* and its conservation across evolution.

## Results

### Replication origins contain two dominating ORC-bound motif sequences that differ in symmetry

To map the genomic binding locations of ORC, we used the spatially resolved method ChEC-seq^26^, where individual fusion of three ORC subunits (Orc1,2,4) to MNase enabled to trigger cleavage of DNA-bound sites through a short calcium pulse. All three ORC subunits localized to the same genomic sites (Fig. S1A), which were consistent with previous ChIP-seq data^27^, as expected (Fig. S1B). We selected Orc1 as our reporter for subsequent analysis.

Orc1 bound to its known motif within a large fraction of replication origins. Our high-resolution data captured its footprint, as described *in-vitro* (45-50 bp^28–32^; Fig. 1B,C) and further enabled the refinement of the motif sequence. This highlighted two variants that dominate strongly bound origins, which we denote as ‘symmetric’ or ‘asymmetric’ (Fig. 1D). Both variants contained a TTTAT stretch at the proximal [5:10] location but differed at the distal [11:16] location, which contained either a stretch of TTTxT, or an TTTAG sequence. Other origins included motif sequences that varied in the [5:10] and [11:16] positions, but those were fewer in count and weakly bound. Finally, Orc1 localized also to loci undergoing transcriptional silencing, where it acted as a silencer^25^. Those included telomeric “X elements”, which contain the ORC motif, but also promoters of sub-telomeric/mating locus silenced genes lacking the motif (Fig. 1E,F). Below we examine the need of various ORC elements for binding these classes of sites.

### A BAH domain explains Orc1 binding and activity at silenced loci

The function of Orc1 in gene silencing depends on its N-terminal BAH domain^38–45^. Early studies reported that this domain is generally required for binding replication origins^46^. We revisited this by mapping the binding profile of a BAH-deleted Orc1 mutant. The BAH-deleted Orc1 was lost from silencing-associated sites, but remained bound to replication origins (Fig. 2A,B). We also tested the binding of the 375-aa long Orc1 N-terminal tail containing the BAH domain (Fig. 2C). While showing no binding at replication origins, this isolated element retained strong binding to all silencing-associated sites which was, again, BAH-dependent (Fig. 2D,E).

**Figure 2.**
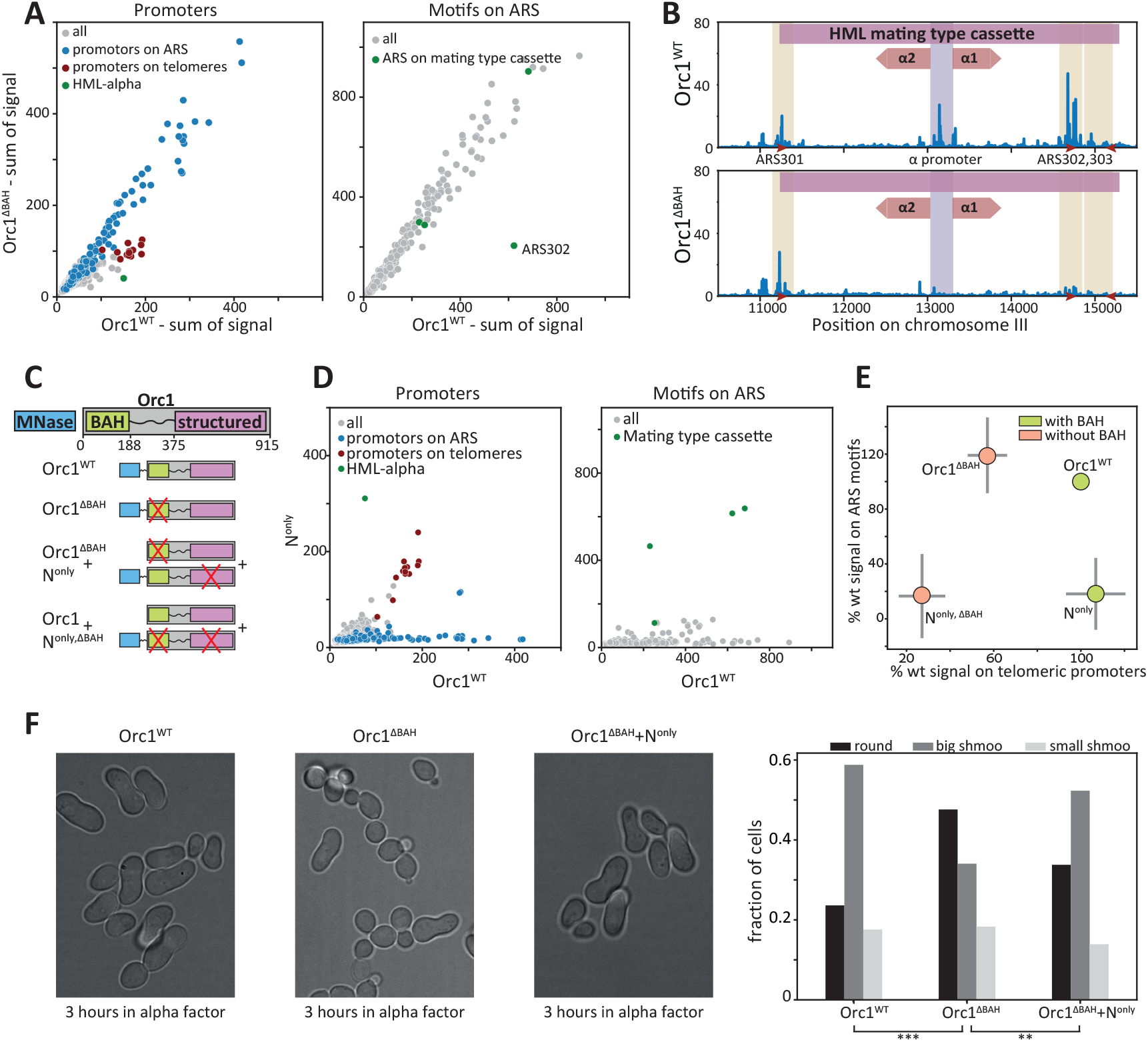
Orc1-BAH domain is necessary and sufficient for Orc1 binding and activity in silenced loci. A-B: *The Orc1 BAH domain is required for binding telomeric promoters but not replication origins:* Shown in A is the average binding on promoters (left) or replication origins (right), comparing the BAH-deleted Orc1 mutant to the full Orc1. Each dot corresponds to one respective sequence, positioned by the overall number of reads assigned to this sequence. Read distribution over one segment – the mating type cassette - is also shown (B). Colors in (A) indicate promoters overlapping ARS elements (blue), promoters of telomere-located silenced genes (red), and promoter of silenced HML-alpha mating-type cassette (green). Colors in (B) indicate mating factor alpha ORFs (red), promoter (gray) and origins (green), while ORC binding motifs are shown as red arrows. C-E: *Orc1 N-terminal tail binds to silenced loci in a BAH-dependent manner:* We examined the binding of an isolated Orc1 N-terminal tail and several of its mutants, as illustrated schematically in (C). Shown are comparisons between the binding of the isolated Orc1-N terminal tail and the full Orc1 on promoters and origins (D, as in A above). The effects of the Orc1-N tail mutants are summarized in (E), showing the average change in binding (%) (with standard deviation in gray) over telomeric promoters lacking the ORC motif (x axis), and origins that do contain the motifs (y-axis). (See Fig. S2 for additional Orc1 N-terminal tail mutants). F. The *Orc1-N terminal tail rescues silencing phenotype of a strain deleted from the Orc1 BAH domain:* A-type haploid cells of the indicated strains were incubated in alpha factor for 3 hours and then visualized under the microscope. >600 cells were segmented and classified into 3 groups: round cells (insensitive to alpha factor), big shmoo (elongated, cell cycle arrested cells) and small shmoo (slightly elongated: ambiguous classification). (Left) examples of cells from each strain, (right) the fraction of cells of each type in the different strains.

To examine whether the Orc1 N-terminal tail is sufficient for gene silencing itself, we quantified cell morphology changes associated with silencing. Specifically, cells of mating type “a” harbor a mating type cassette whose silencing is required for cell elongation upon exposure to alpha factor (“shmooing”). Loss of silencing at the mating-type cassette abolishes this response and this was indeed reported for BAH-deleted deleted cells^47^. As predicted by the binding profile, the isolated Orc1 N-terminal tail was sufficient to rescue this silencing phenotype (Fig. 2F). We conclude that the BAH domain explains Orc1 binding and activity at silencing-associated sites.

### The Orc4 insertion helix is required for binding at asymmetric, but not symmetric motif sequence

The Orc4 insertion helix (Orc4-IH) was described in the cryo-EM structure as the only base-contacting ORC region in the DNA major groove. Subsequent studies raised the hypothesis that Orc4-IH determines motif specificity^23,24^, and showed that Orc4-IH mutants exhibit slow growth, along with perturbed origin firing and replication-helicase loading^23,24^.

We revisited the role of Orc4-IH by directly mapping the genomic binding of Orc4 mutated at two base-contacting residues^22^(substituting F485 and Y486 to Alanine; Fig. 3A,S3). This mutant showed reduced binding at some origins (e.g. ARS1528, Fig. 3B) but, perhaps unexpectedly, retained strong binding at others, including the origin used for structure determination (ARS305; Fig. 3B). The binding effects distinguished between the two motif sequences, with the asymmetric motif exhibiting a much higher loss in binding (Fig. 3C,D). This was also the case in telomeres, where motif-lacking silencing sites or symmetric motifs were not affected, while asymmetric motif sites showed high loss of binding (Fig. 3E). Note that Orc4-IH contacts nucleotides within the second T-stretch distinguishing the symmetric and asymmetric motifs. We conclude that Orc4-IH expands ORC binding to sequences of reduced symmetry and T-content.

**Figure 3.**
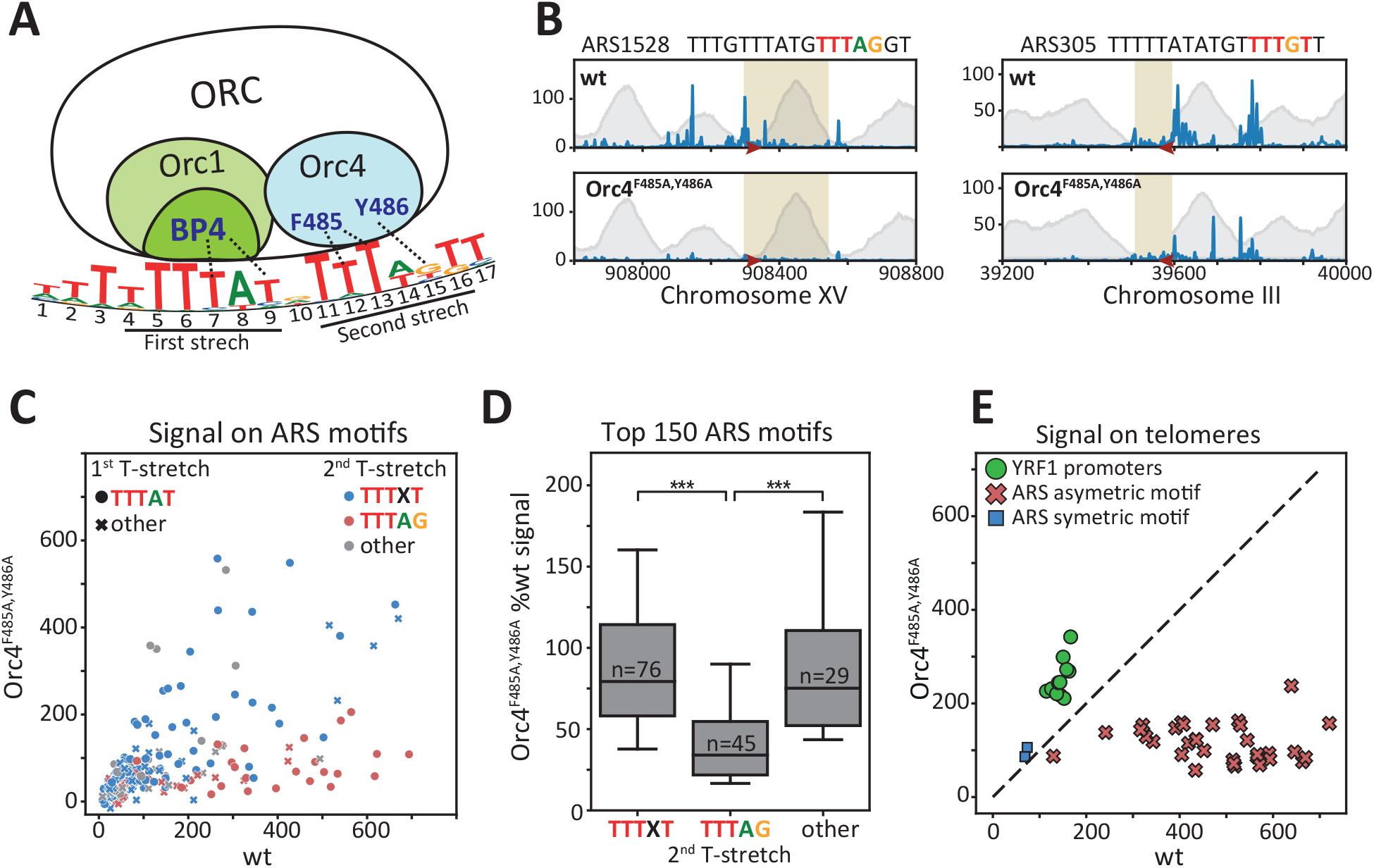
Orc4-IH stabilizes ORC binding specifically in replication origins containing asymmetric motif variant. *A. The two ORC components directly contacting the ORC motif:* a schematic representation based on the cryo-EM structure^22^. *B. Orc4-IH is required for ORC binding at a subset of origins:* Shown is the Orc1 binding profiles at two representative replication origins (ARS1528 and ARS305), comparing binding of wild-type cells (wt) and cells carrying Orc4-IH mutation (Orc4-F485A,Y486A). Annotations as in Fig. 1B above. C-D: *Orc4-IH mutation abolishes binding specifically at origins carrying the asymmetric motif:* A comparison of Orc1 binding signal across all origins is shown in (C), as measured in Orc4-F485A,Y486A mutant vs. wild-type cells. Each dot represents an origin, and its shape and color indicate the associated motif variant, as specified. Also shown (D) is the distribution of binding changes across origins containing the indicated motif variants. Asterisks denote p-values: *p<0.01, **p<0.001, p***<0.0001. *E. Orc1 retains strong binding at silenced loci in Orc4-IH mutant:* Orc1 binding in the Orc4-F485A,Y486A mutant vs. the wild-type cells on the indicated telomeric elements.

### Orc1-BP4 is required for binding at replication origins, but is dispensable at silenced loci

The finding that Orc4-IH is required for binding at the asymmetric, but not symmetric motif sequence, motivated us to examine Orc1-BP4, the second base-interacting element seen in the cryo-EM structure. Previous studies using structural modeling^48,49^ and conservation analysis^50^ predicted a role for this BP in origin binding (Fig. 4A), and mutating the *Drosophila* Orc1-BP reduced ORC-DNA affinity *in-vitro*^*32*^. Orc1-BP4 is essential for growth, which supports its general role in binding, but also complicates its analysis.

**Figure 4.**
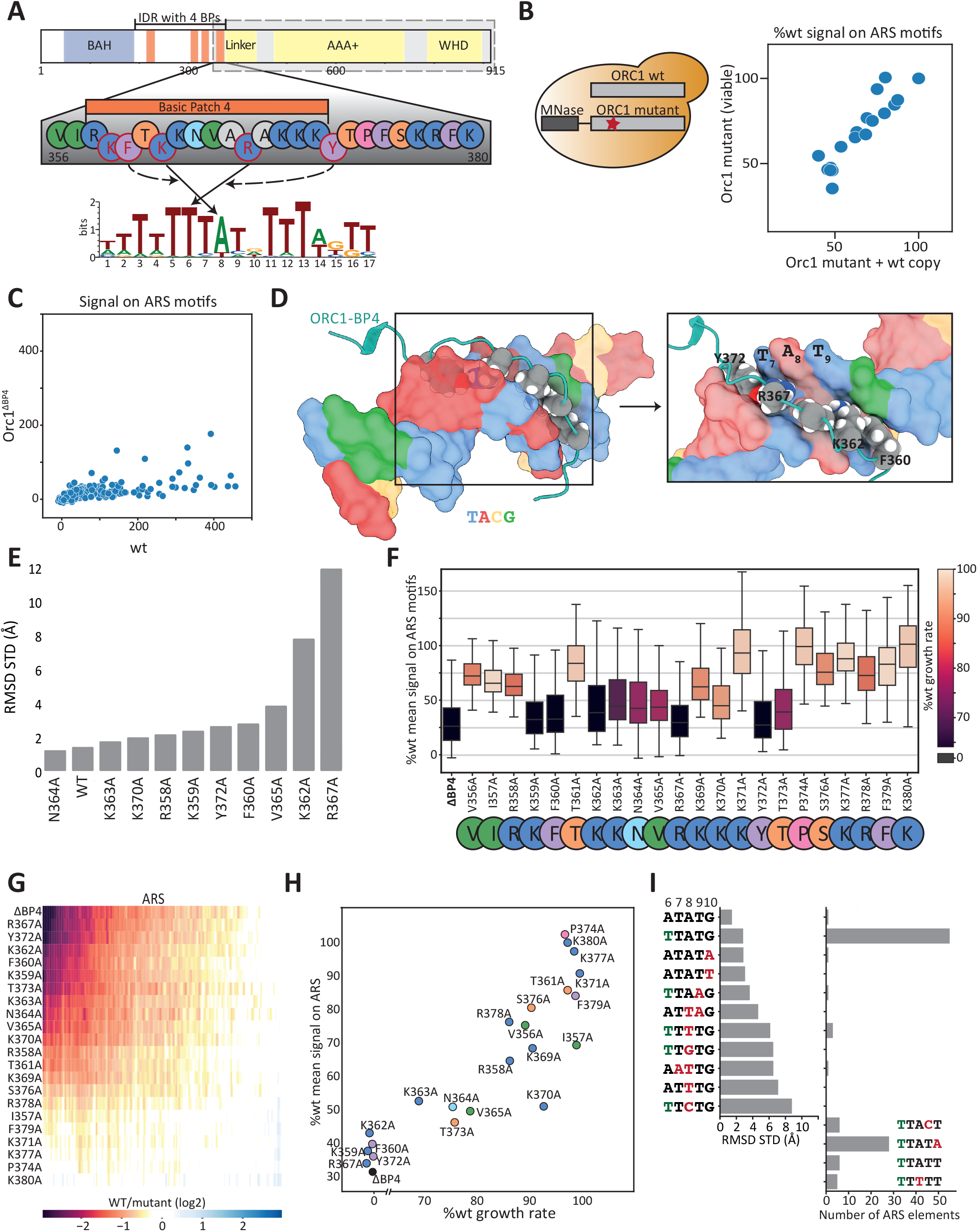
Orc1-BP4 stabilizes binding at all replication origins, depending on multiple residues beside the direct base-contacting ones. *A. Orc1-BP4 interaction with the ORC motif:* A scheme based on the cryo-EM structure^22^ Full and dashed arrows indicate specific DNA-base contacts and predicted “supportive” residues, respectively. *B. Binding profile of lethal mutantations is measured by adding an exogenous wild-type allele:* Deleting Orc1-BP4 or mutating its residues often resulted in a lethal or slow-growth phenotype (see below). To still map their binding, we engineered a strain containing an intact Orc1 copy and a mutant Orc1 fused to MNase, as illustrated (left). This enabled mapping of the mutant allele, as we verified by profiling viable mutants in the presence or absence of the wild-type copy and subsequently comparing them. Shown here are the median binding signal across origins in the presence or absence of the additional wild-type Orc1 copy (right) (see Fig. S4A for specific mutation comparisons). *C. Orc1-BP4 deletion abolishes Orc1 binding at all origins:* Shown is the binding signal of Orc1 lacking BP4 (y-axis) compared to intact Orc1 (x-axis) across all replication origins. *D. Atomic view of Orc1-BP4 (cyan) inserted into the DNA minor groove, as described by the cryo-EM model*^*22*^: Direct binding amino acids located in the minor groove are labeled, and each nucleotide is color-coded: T in blue, A in red, C in orange, and G in green. *E. MD simulations predictions of Orc1-BP4 mutant effects:* Shown is the predicted conformational stability of the indicated BP4 mutants. Conformational stability was estimated by the plasticity of Orc1-BP4 localization at its binding site, as quantified by the standard deviation of the root mean square deviation (over all the simulation steps) relative to the localization seen in the cryo-EM structure (PDB ID 5ZR1). *F-H: Multiple BP4 residues beyond the base-contacting ones contribute to motif binding:* Shown in (F) are the distributions of binding signals of the indicated Orc1-BP4 mutants, relative to the wild-type Orc1-BP4, across all origins. Colors indicate mutant growth rates. Also shown are the fold-change in binding relative to wild-type BP4 across individual origins (G), and a direct comparison of the average change in binding and mutant growth rate (H). *I. MD simulations do not explain the conservation of the Orc1 binding motif:* Shown on the left are conformational heterogeneities observed when simulating binding of intact Orc1-BP4 to the indicated motif variants. Mutations are marked in red, while T6 of the conserved motif is denoted in green. Results are compared to the number of occurrences of the specified variants among the 150 top-bound origins in the genome (right).

To overcome this, we mapped binding of Orc1-BP4 mutants, while simultaneously introducing an exogenous (untagged) Orc1 copy. Verifying that this exogenous copy does not alter the measured binding profile of non-essential mutations (Fig. 4B,S4A), we proceeded to analyze the binding of BP4-deleted Orc1. Deleting BP4 dramatically reduced Orc1 binding at most replication origins (Fig. 4C), consistent with its insertion within the proximal T-stretch^50^ common to both motif sequences. Binding at silencing-associated sites was not affected (Fig. S4B).

Orc1-BP4 binding to DNA differs from the common paradigm of DNA-protein binding, as it involves the insertion of a disordered region within the minor groove (Fig. 4D), providing little specificity. We were therefore interested in exploring the properties of this binding, and for this we used molecular dynamics (MD) simulations. We initiated the simulation with Orc1-BP4 being at the stable DNA-bound configuration, as described in the cryo-EM structure^22^ (PDB ID: 5ZR1), and measured its subsequent confinement, as indicated by the standard deviation of the root mean square deviation (RMSD) across all the simulation steps. The intact BP4 showed little movement, as expected, but this confinement was lost when mutating either of the base-contacting residues to alanine (K362A and R367A), validating our simulations (Fig. 4E). Testing eight additional mutants at different BP4 locations, we found that all retained DNA confinement. We conclude that local BP4-base interactions are sufficient to explain its DNA binding, while the surrounding residues are of smaller apparent effects.

We also tested these mutation effects experimentally by performing an alanine scan covering the 23 non-alanine residues within and surrounding Orc1-BP4 (Fig. 4A). Contrasting our MD simulations, 17 out of 23 replacements did cause a substantial reduction in DNA binding (Fig. 4F). Effects were correlated across mutants (Fig. 4G) with PCA analysis attributing 88% of the variation to a single component (Fig. S4C), and were well explained by the distance to the two base-contacting residues (K362, R367) or their minor-groove embedded assisting residues (F360, Y372) (Fig. 4D). We also measured growth phenotypes of those same mutants, using strains that lacked the endogenous copy, and those correlated well with the loss of binding (R^2^=0.90, p value=7.5*10-9), including the apparent essentiality (inability to generate) of the five mutants showing strongest binding effects (K362A, R367A, F360A, Y372A, K359A) (Fig. 4F,H). Therefore, the *in-vivo* Orc1-BP4-DNA binding depends on multiple BP4 residues, surrounding the base-contacting ones.

The partial agreement between the MD simulations and the measured phenotypes of BP4 mutants suggests that additional factors, not included in the simulations, influence BP4-DNA binding *in-vivo*. BP4, for example, could interact with other ORC components, and we noted some indications for this within the structure (Fig. S4D). Perturbation of such intramolecular interactions within ORC1 may affect its DNA-binding affinity. Alternatively, some mutations might affect transient BP4 interactions with nonspecific DNA sites that are required for guiding Orc1 to its high-affinity binding at the specific site. Such mutations may affect the overall binding preferences of ORC1 along genomic DNA but would be invisible to our MD simulations or the structure. In this latter case, MD predictions would also be blind to mutating DNA bases participating in those transient interactions because quantifying their effect demands dynamics and kinetic characterization^51^ which is not accessible to the atomistic modeling.

To test this, we simulated BP4 interaction with different mutated DNA sequences (Fig. 4I). First, we noted that the motif sequence used in the structure (PDB ID: 5ZR1), and therefore our simulations, differed from the canonical one, although it was still strongly bound in-vivo. Mutating this sequence to retrieve the canonical abundant motif sequence retained strong binding, while mutating the conserved A base at the center of BP4-DNA interactions (position 8 in the motif) abrogated BP4 confinement, validating our simulations. Notably, mutating other conserved residues 2bp apart from the core interactions, had no or little effect on confinement within our simulations, although the respective DNA sequences were lacking, or weakly bound *in-vivo* (Fig. 4I). Taken together, those results suggest that transient BP4 interactions with semi-specific DNA sites guide ORC to its high-affinity binding, but these are invisible in both the MD simulations and structural analysis, which report on the local binding stability.

### BP3 is essential for ORC binding in the absence of Orc4-IH interacting residues

Our results so far have revealed that binding at the asymmetric motif sequence requires both Orc1-BP4 and Orc4-IH, while binding the symmetric motif requires only Orc1-BP4. This raised the question of whether the later binding depends on an additional ORC element. Since Orc1-BP4 and Orc4-IH were the only base-contacting elements seen in the structure, we next considered the Orc1 N-terminal tail that was absent from the Orc1 used for the structure analysis^22^(Fig. 5A). This tail borders BP4 and includes three additional basic patches. We found that truncating the proximal two had no detectable effect on ARS binding, but a longer truncation that included BP3 reduced ORC binding (Fig. 5B). We further profiled four additional Orc1 mutants: one that lacked only BP3 (residues 320-340), two that lacked BP4 (residues 357-375 or residues 340-375), and a mutant that lacked both BP3 and BP4 (residues 320-375). Consistently, deleting BP3 alone led to a moderate reduction in binding, while co-deletion of BP3 and BP4 accentuated the effect of BP4 deletion (Fig. 5C,D).

**Figure 5.**
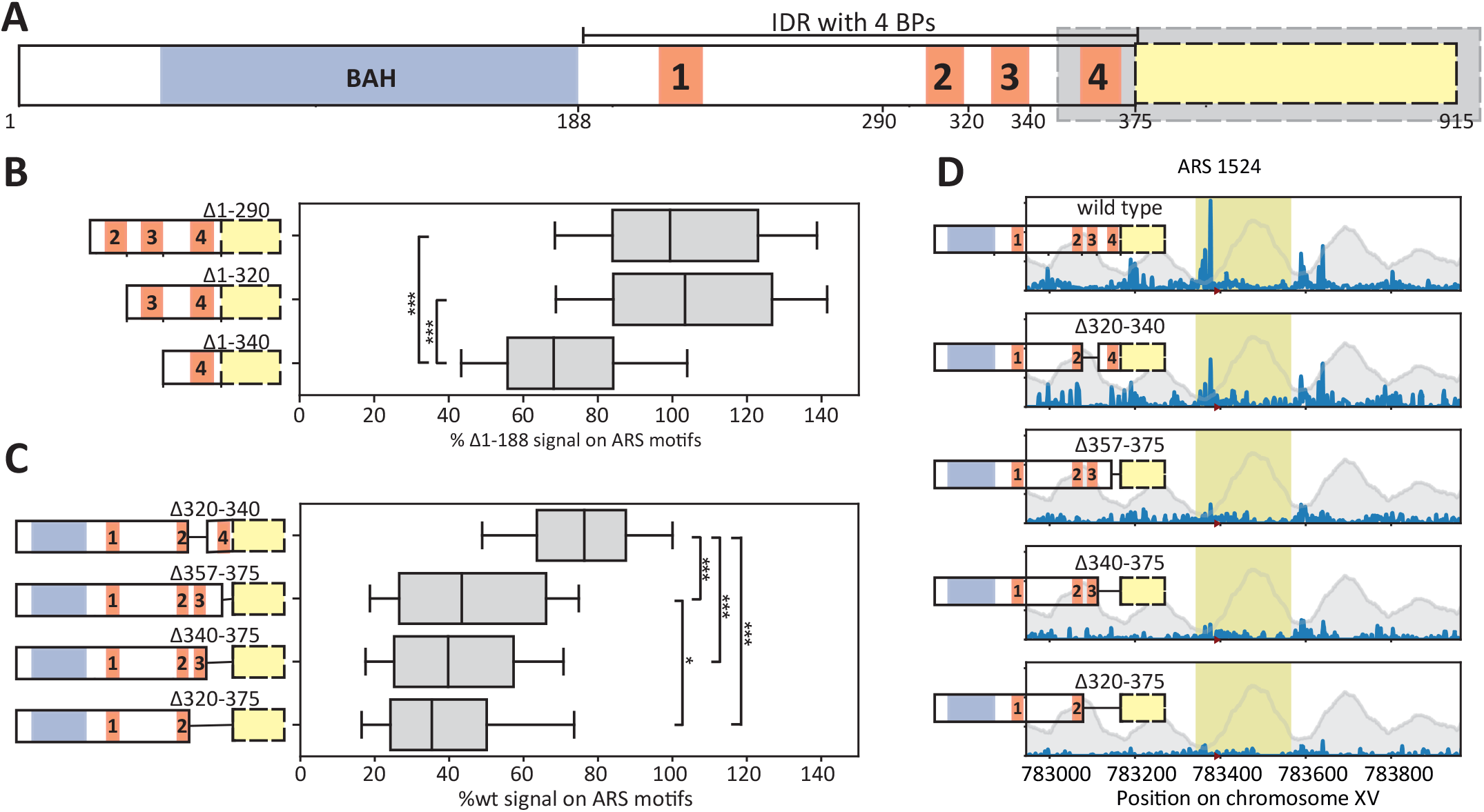
ORC1-BP3 contributes to origin binding. A. A. *Schematic representation of Orc1 N-terminal tail*: Colors as in Fig. 1A. Note the four basic patches (BP4) within the intrinsically disordered region (IDR). The cryo-EM structure^22^ was solved using a truncated Orc1, containing the 355-914 AA (Gray). B-D: *Orc1-BP3 deletions reduces origin binding:* Shown in (B, C) are the distributions of origin binding of the indicated mutants. Data are shown relative to the 188aa-truncated strain (BAH deletion, B) or intact Orc1 (C). The respective reads distribution across a selected origin is also shown (D).

As expected, Orc1-BP3 deletion affected the symmetric motif sequence (Fig. 6A). We therefore hypothesized that ORC1-BP3 and ORC4-IH play a complementary role in stabilizing ORC and examined this by combining the Orc1-BP3 and Orc4-IH mutations (Orc4 F485A and Y486A). Indeed, while the individual mutants were viable, we could not generate the combination, suggesting lethality. To verify this interaction at the level of ORC binding, we introduced the Orc4-IH mutations into cells carrying a deleted Orc1-BP3 tagged with MNase, as well as an intact (unlabeled) Orc1 (as illustrated in Fig. 4B). As predicted, BP3 was essential for ORC binding at replication origins when Orc4-IH was mutated (Fig.6A,C), with binding remaining only at silencing-associated loci (Fig.6D). We conclude that Orc1-BP3 and Orc4-IH can both complement Orc1-BP4 in stabilizing ORC binding. In this, Orc4-IH is required for the asymmetric motif sequence, while Orc1-BP3 can bind the symmetric one.

**Figure 6.**
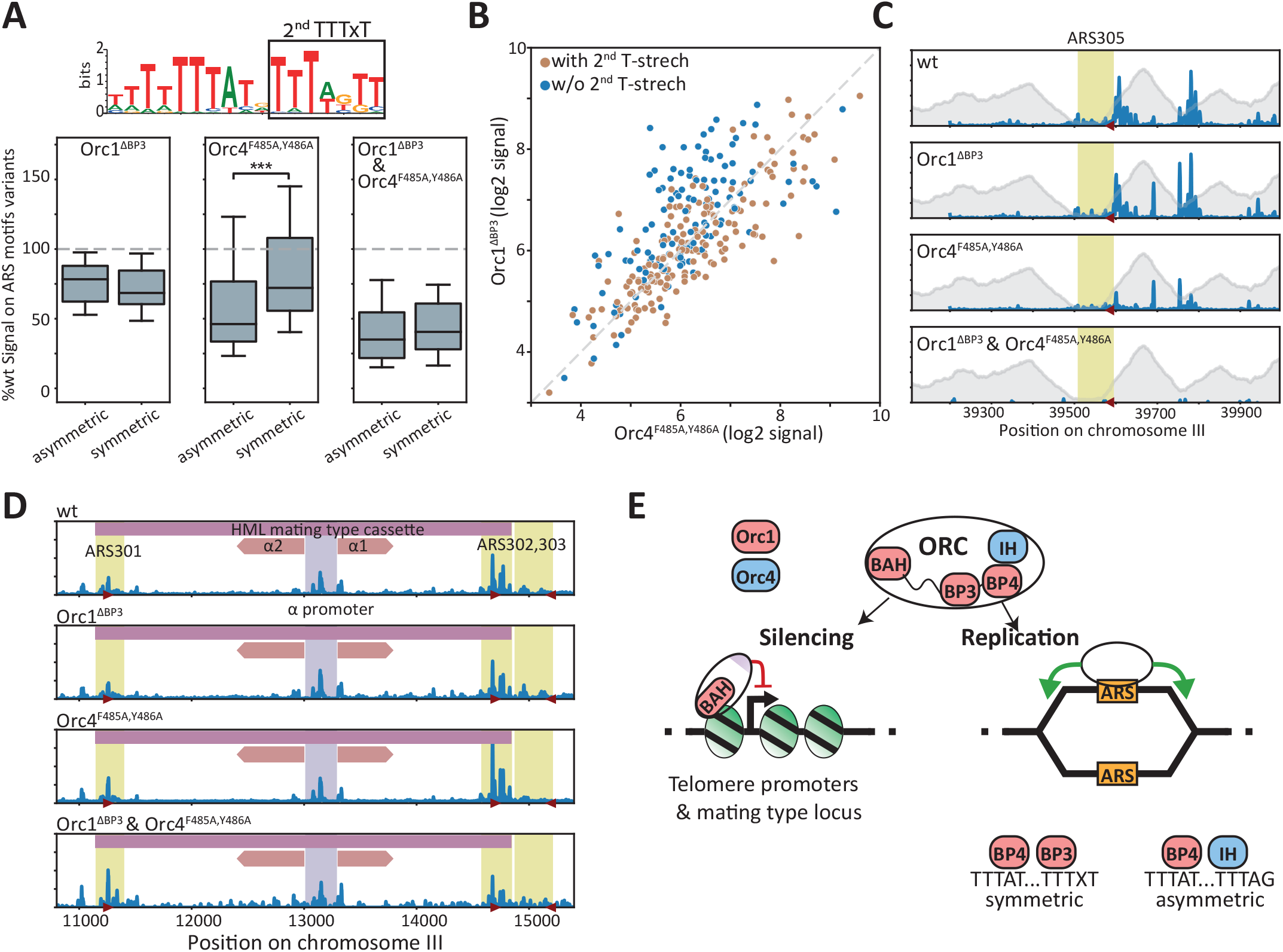
Orc1-BP3 and Orc4-IH play complementary roles in stabilizing ORC binding at replication origins. A-C: *Orc1-BP3 is essential for origin binding in Orc4-IH mutant cells:* Shown in (A) are the distributions of origin binding in the indicated single and double mutants. Origins containing the symmetric or asymmetric motifs variants are shown separately. Also shown are the effects of the individual mutants on individual origins, with colors indicating the associated motif variant (B), and the reads aligned to the origin in the cryo-EM structure^22^(C, annotation as in Fig. 1B). D. Distributions of binding of the indicated mutants to the silenced silenced HML mating type cassette. E. *A model – three distinct modes of ORC-DNA binding:* In promoters of silenced genes lacking the motif, the Orc1-BAH domain is necessary and sufficient for binding. In replication origins, ORC binding differs depending on the motif variants. Orc1-BP4 is required for both sets, but an additional anchor is required to stabilize binding, and this is provided by either Orc4-IH or Orc1-BP3, with the latter able to only supplement Orc4-IH at the symmetric motif variant.

## Discussion

The ORC lacks an apparent DNA binding domain but still localize specifically to certain DNA sequences, which, in budding yeast, are marked by a 17-bp T-rich motif. Structural analysis partially explained this preference by revealing two ORC elements that directly contact five motif bases. One of these (Orc4-IH) inserts within the major groove, while the second (Orc1-BP4) inserts into the minor groove, providing little base discrimination. Those limited interactions raised the question of whether they are sufficient to explain ORC-DNA binding *in-vivo*, motivating our study.

We improved the spatial resolution of ORC binding locations using ChEC-seq, which revealed previously unrecognized classes of ORC binding motifs: a symmetric sequence that contained two T-stretches (a proximal TTTAT and a distal TTTxT) and an asymmetric class that contained the same proximal stretch, but a distal TTTAG stretch. Further analysis revealed that ORC binding differs between those variants. Orc1-BP4 was required for binding at both, consistent with its insertion at the common proximal stretch, while Orc4-IH was only needed for binding to the asymmetric motif. At the symmetric motif, Orc4-IH was not required, and could be compensated by a second basic-patch, Orc1-BP3, that was absent from the structure.

To gain further insights into ORC-DNA binding, we delved deeper into Orc1-BP4 interactions through mutation analysis, combining MD simulations and experiments. It is notable that our simulations reproduced well the phenotypes of the direct contact sites, on both DNA and the peptides. However, they were blind to the effects of additional Orc1 residues or DNA elements positioned a few residues apart. These differences could result from additional interactions not included in our simulations. Yet, their prevalence on both DNA and BP4 itself leads us to favor an alternative explanation, in which ORC-DNA binding depends on transient BP4-motif interactions. Although transient only, these interactions can still dominate the mechanism and speed for reaching the high-affinity site visible to the structure and MD simulations. Modulating the rate of binding to the DNA motif can directly affect the efficiency of high-affinity binding, and therefore functionality^52^.

Based on this, and the revealed symmetry of both the motif and the associated Orc1-BP interactors, we propose a model (Fig. 6E) in which BPs within the Orc1-N terminal tail favor sliding within the minor grooves of T-stretches. This sliding is stabilized when reaching a non-T residue, as in TTTxT. We further propose that two such interactions are required, as individual ones are not strong enough. In motifs that lack the second TTTxT stretch, this role is overtaken by Orc4-IH which interacts with an alternative TTTAG sequence. Further studies are required to test this model.

The *S. cerevisiae* ORC used in this study is unique in preferring a specific sequence motif. Other eukaryotic ORCs do not show this preference, but still localize to specific T-rich genomic locations. Based on the conservation of BPs within the Orc1 N-terminus, we propose that ORC-DNA binding is guided by those BPs through a conserved mechanism common also to *S. cerevisiae*, potentially with slight alterations tuned for the requirements of each genome, as exemplified by Orc4-IH in budding yeast.

## Methods

### Yeast strains

All strains used in this study are derived from the wild-type *Saccharomyces cerevisiae* strain, BY4741 (genotype: MATa his3D1 leu2D0 met15D0 ura3D0). Specific genotypes of all strains used in this study are available in Table S1.

### Genetic manipulations

Yeast strains were freshly thawed before experiments from a frozen stock, plated on YPD plates, and incubated in 30°C. Single colonies were picked and grown in liquid YPD medium at 30°C with constant shaking. DNA editing was generated with the CRISP-Cas9 system^53^. To this end, a PCR-amplified repair template or a synthetic oligo harboring the desired modification flaked by 50-bp homology was co-transformed alongside the bRA89 plasmid bearing Cas9 and the locus-specific 20-bp guide-RNA (from James Haber, Addgene plasmid no. 100950). Ligation of the locus-specific guide-RNA into the bRA89 plasmid was performed for each guide-RNA as previously described^54^. After validation of yeast positive colonies, the bRA89 plasmid was lost by growth in YPD, followed by the selection of colonies that lost the bRA89 Hygromycin resistance.

All transformations were performed using the standard LiAc/SS DNA/PEG method^55^. Briefly, a single colony was inoculated in fresh liquid YPD, grown to saturation overnight, diluted into fresh 5 ml YPD, and grown to OD_600_ of 0.5. The cells were then washed with DDW and then with LiAc 100 mM, and resuspended in transformation mix (33% PEG-3350, 100 mM LiAc, single stranded salmon sperm DNA, plasmid and the DNA repair). The cells were incubated at 30°C for 30 minutes followed by a 30-minute heat shock at 42°C. The cells were then plated on YPD plates and grown overnight in 30°C for recovery.

For strains containing an additional copy of the wild-type Orc1, a synthetic DNA block coding the wild-type ORC1 protein with numerous synonymous mutations (sequence in supp Document S1) was inserted into the genome. This was done along with the original Orc1 promoter and into the Bar1 locus, creating a bar1 deletion. Additional genetic modifications were carried out by specifically targeting only the endogenous Orc1, using guide-RNAs that did not match the synonymous sequence of the synthetic Orc1.

### Chec-seq experiments

The experiments were performed as described previously^26^, with some modifications. Yeast strains were freshly thawed before experiments from frozen stocks, plated on YPD plates, and incubated at 30°C. Single colonies were picked and grown overnight at 30°C with constant shaking in liquid YPD medium to stationary phase. Then, the cultures were diluted into 15 mL fresh YPD media and grown overnight at 30°C to reach log-phase in the following morning. Cultures were pelleted at 1500 g for 2 min and resuspended in 0.5 mL buffer A (15 mM Tris pH 7.5, 80 mM KCl, 0.1 mM EGTA, 0.2 mM spermine, 0.5 mM spermidine, 13 cOmplete EDTA-free protease inhibitors (one tablet per 50 mL buffer), 1 mM PMSF) and then transferred to 2.2 mL 96-well plates (LifeGene). Cells were washed twice in 1 mL Buffer A. Next, the cells were resuspended in 150 mL Buffer A containing 0.1% digitonin, transferred to an Eppendorf 96-well plate (Eppendorf 951020401), and incubated at 30°C for 5 min for permeabilization. Next, CaCl_2_ was added to a final concentration of 2 mM to activate the MNase and incubated for exactly 30 seconds. The MNase cleavage was stopped by adding an equal volume of stop buffer (400 mM NaCl, 20 mM EDTA, 4 mM EGTA, and 1% SDS) to the cell suspension. After this, the cells were treated with Proteinase K (0.5 mg/mL) at 55°C for 30 minutes. An equal volume of phenol-chloroform pH=8 (Sigma-Aldrich) was added, vigorously vortexed, and centrifuged at 17,000g for 10 minutes to extract DNA. After phenol-chloroform extraction of nucleic acids, the DNA was precipitated with 2.5 volumes of cold 96% EtOH, 45 mg Glycoblue (Thermo Fisher), and 20 mM sodium acetate at –80°C for >1 hr. DNA was centrifuged (17,000 g, 4°C for 10 min), the supernatant removed, and the DNA pellet washed with 70% EtOH at room temperature. DNA pellets were dried and resuspended in 30 mL RNase A solution (0.33 mg/mL RNase A in Tris-EDTA [TE] buffer [10 mM Tris and 1 mM EDTA]) and treated at 37°C for 20 min. To enrich for small DNA fragments, the samples went through reverse SPRI cleanup by adding 0.83 SPRI beads (Ampure XP). The supernatant was collected, and the remaining small DNA fragments were purified by adding an additional 13 SPRI beads and 5.43 isopropanol, and incubating for 5 min at RT. Beads were washed twice with 85% EtOH, and small fragments were eluted in 30 mL of 0.13 TE buffer.

### Next Generation Sequencing library preparation

Library preparation was performed as described by Skene and Henikoff^56^ with slight modifications. DNA fragments after RNase treatment and reverse SPRI cleanup served as an input to end-repair and an A-tailing (ERA) reaction. For each sample, 20 mL ERA reaction (13 T4 DNA ligase buffer [NEB], 0.5 mM dNTPs, 0.25 mM ATP, 2.75% PEG 4000, 6U T4 PNK [NEB], 0.5U T4 DNA Polymerase [Thermo Scientific] and 0.5U Taq DNA polymerase [Bioline]) was prepared and incubated for 20 min at 12°C, 15 min at 37°C and 45 min at 58°C in a thermocycler. After ERA reaction, reverse SPRI cleanup was performed by adding 0.53 (10 mL) SPRI beads (Ampure XP). The supernatant was collected, and the remaining small DNA fragments were purified with additional 1.33 (26 mL) SPRI beads and 5.43 (108 mL) isopropanol. After washing with 85% EtOH, small fragments were eluted in 17 mL of 0.13 TE buffer; 16.4 mL elution was taken into 40 mL ligation reaction (13 Quick ligase buffer [NEB], 4000U Quick ligase [NEB], and 6.4 nM Y-shaped barcode adaptors with T-overhang^57^ and incubated for 15 min at 20°C in a thermocycler. After incubation, the ligation reaction was cleaned by performing a double SPRI cleanup: first, a regular 1.23 (48 mL) SPRI cleanup was performed and eluted in 30 mL 0.13 TE buffer. Then and instead of separating the beads, an additional SPRI cleanup was performed by adding 1.33 (39 mL) HXN buffer (2.5 M NaCl, 20% PEG 8000) and final elution in 24 mL 0.13 TE buffer; 23 mL elution were taken into 50 mL enrichment PCR reaction (13 Kappa HIFI [Roche], 0.32 mM barcoded Fwd primer and 0.32 mM barcoded Rev primer^57^, and incubated for 45 s in 98°C, 16 cycles of 15 s in 98°C and 15 s in 60°C, and a final elongation step of 1 min at 72°C in a thermocycler. The final libraries were cleaned by a regular 1.13 (55 mL) SPRI cleanup and eluted in 15 mL 0.13 TE buffer. Library concentration and size distribution were quantified by Qubit (Thermo Scientific) and TapeStation (Agilent), respectively. For multiplexed next-generation sequencing (NGS), all barcoded libraries were pooled in equal amounts, the final pool diluted to 2 nM and sequenced on NovaSeq 6000 (Illumina).

### Process and analysis of ChEC-seq data

Raw paired end reads from ChEC-seq libraries were demultiplexed using bcl2fastq (Illumina), and adaptor dimers and short reads were filtered out using cutadapt^58^ with parameters: “-O 10 –pair-filter=any -max-n 0.8 - action=mask”. Filtered reads were subsequently aligned to the *S. cerevisiae* genome R64-1-1 using Bowtie 2 ^59^ with the options “-end-to-end -trim-to 30 -very-sensitive”. The genome coverage of fully aligned read pairs was calculated using the fragment ends which correspond to the actual MNase cutting sites. Each sample was normalized to one million reads to control for sequencing depth reads. ARS, ORF and telomere annotations were taken from the SGD project^60^. For each ARS element, the sub-sequence within the defined ARS borders, with the highest motif score was selected, and the signal within a window of 75 pb upstream and downstream to it was calculated. Motif scores were calculated using the PWM matrix from YeTFaSCo^37^.

### Automated growth rate experiments

Cells were grown to stationary phase in YPD media in 96-well plates under constant shaking and at 30°C. Then, cells were diluted with fresh YPD media to OD_600_ 0.1. Plates were inserted into an automated handling robot (EVOware, Tecan Inc.) in which cells were grown in an incubator under constant shaking and 30°C. The robot was programed to take the plates out of the incubator every 30 minutes, vortex the plates, and measure the OD (using infinite200 reader, Tecan Inc.). Experiments lasted for approximately 40 hours. OD measurements were parsed and processed from the reader output files. Growth rates were calculated as the maximal slope of the OD measurements (versus time). For each strain the mean value over 6 repeats was calculated.

### Alpha factor imaging

Yeast strains (with bar1 deletion) were freshly thawed before experiments from frozen stocks, plated on YPD plates, and incubated at 30°C. Single colonies were picked and grown overnight at 30°C and constant shaking in liquid YPD medium to stationary phase. Then, the cultures were diluted into fresh YPD media and grown overnight at 30°C to reach log-phase in the following morning (OD_600_ 0.2). α-factor was added to the cells to a final concentration of 5 ng/ml α-factor (GenScript RP01002) for 3 hours. Cells were attached to 18-well μ-Slide (Ibidi) pre-treated with Concanavalin A (ConA): 75 μL of 0.2 mg/μL ConA (Sigma-Aldrich) was added to each well and incubated at room temperature for 15 minutes. Subsequently, liquid was removed, and wells were air dried for 30 minutes. Then, 100 μL of the cell samples were added to the wells and incubated for 15 minutes, followed by a gentle wash with fresh media (with α-factor) to remove unattached cells. Imaging was done using a DMi8 microscope from Leica, integrated into a Dragonfly 505 from AndOr. AndOr’s Fusion software was used for data acquisition.

### Process and analysis of imaging data

Cells were segmented using a freely available yeast segmentation software^61^. Cell classification was performed semi-manually by presenting a single random cell each time from all images, without disclosing the specific strain, for the user to label.

### All-atom molecular dynamics simulations

The simulations were performed using GROMACS^62^. The initial conformation was the solved structure of the ORC-DNA complex^22^ (PDB ID 5ZR1). The simulations included the Orc1-BP4 region and its surrounding residues (Orc1 aa 356-373), along with a subset of 20 nucleotides of each DNA strand that include the ARS consensus sequence (TGGTTTTTATATGTTTTGTT). For each mutation (in the protein or DNA), 3 repeats were done, each lasting 500 steps. The root mean square deviation (RMSD) of all atoms relative to the localization seen in the cryo-EM structure was calculated for each step in the simulation. The standard deviation of all RMSD values of a simulation was calculated, and averaged over all repeats.

### Box plots

Boxes capture 25 to 75 percentiles; median is shown inside the boxes and the whiskers capture 10 to 90 percentiles. P-values are shown in asterisks: * smaller than 0.05, ** smaller than 0.01, *** smaller than 0.001.

## Acknowledgments

We thank Joseph Steinberger for his guidance and assistance in conducting the microscopy experiments. We thank Gilad Yaakov, Sagie Brodsky, Felix Jonas and Alon Chappleboim for their insightful comments and suggestions on the manuscript. Special thanks are also due to the entire Barkai lab for creating a stimulating atmosphere and engaging in fruitful discussions. The funding for this work was provided by the European Research Council (ERC).

