## Supplemental Information for "Ordered and disordered regions of the Origin Recognition Complex (ORC) combine to direct in-vivo binding at symmetric and non-symmetric motif sequences"

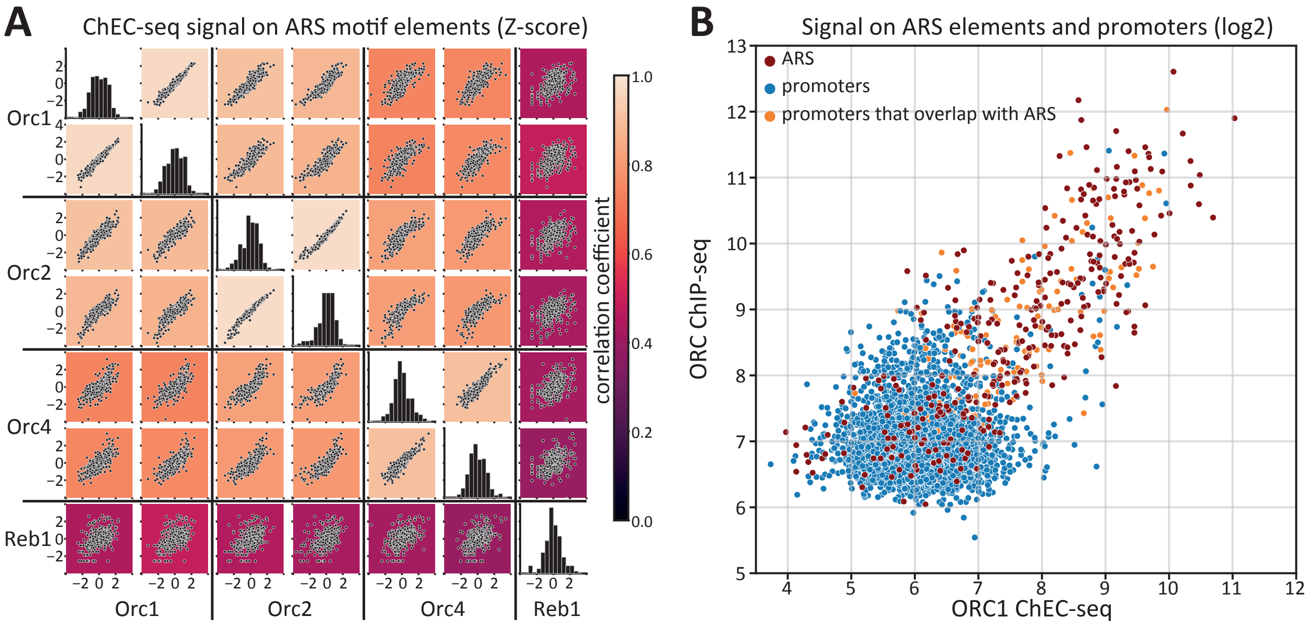


**Figure S1. Localization of ORC ChEC-seq data at ARS elements and its correlation with prior datasets.**

A. ChEC-seq experiments were conducted on strains where MNase was fused to Orc1, Orc2, Orc4, or Reb1 (for comparative analysis). Two representative repeats for each Orc protein are shown. Each point denotes the cumulative signal (z-score) surrounding the ARS element consensus sequence. Colors depict the correlation coefficient.

B. Comparison between Orc1 ChEC-seq signal and ORC ChIP-seq signal (data from Belsky et al.[1]) at ARS elements and promoters as specified.

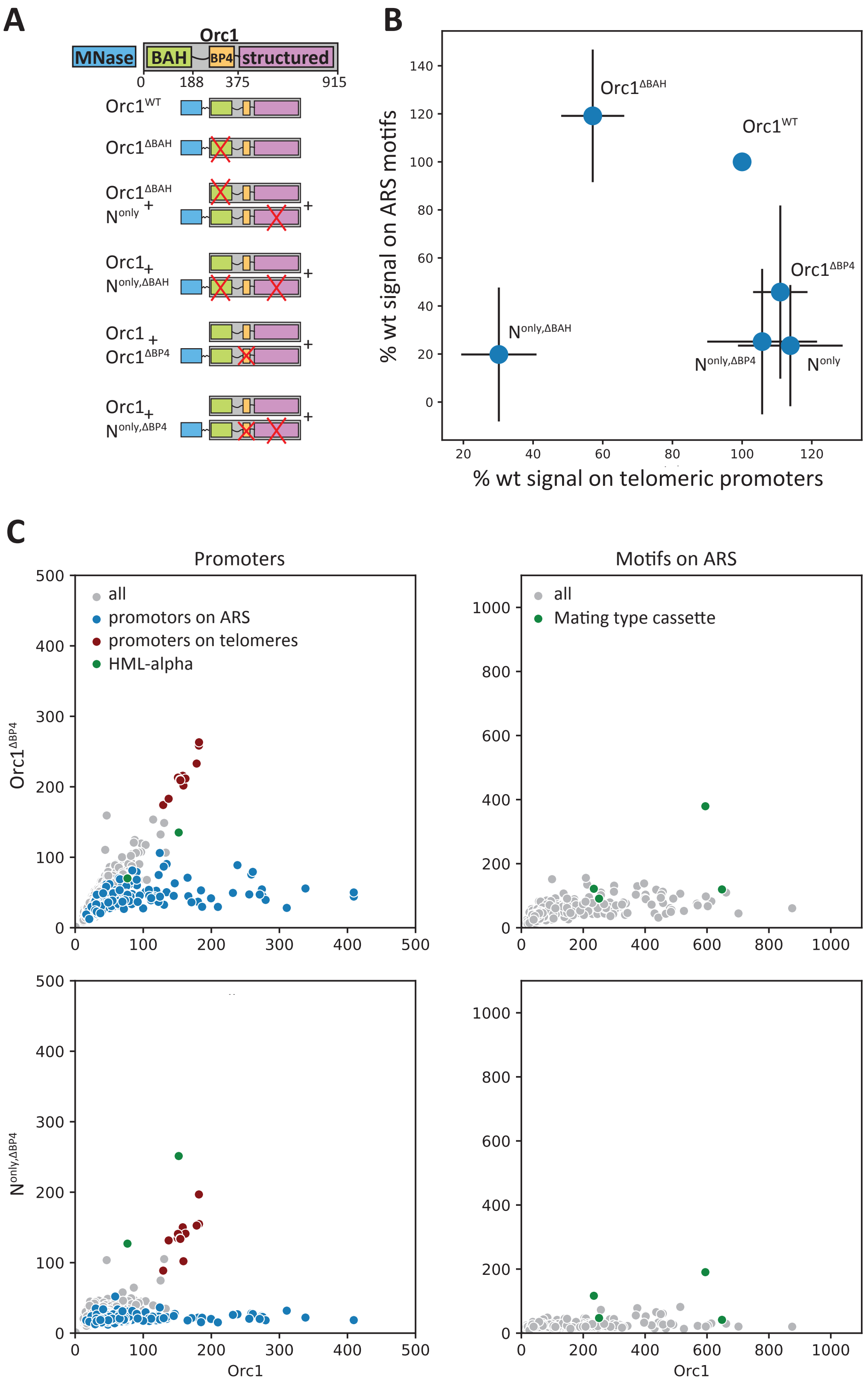


**Figure S2.** **Orc1 N-terminal tail binds to silenced loci in a BAH-dependent manner.**
All the mutants that we have tested for the isolated Orc1 N-terminal tail are illustrated schematically in (A). The effects of the Orc1-N tail mutants are summarized in (B) , showing the average change in binding (%) (and standard deviation in gray) over telomeric promoters lacking the ORC motif (x axis), and origins that do contain the motifs (y-axis). Orc1-BP4, which is necessary for binding to ARS motifs (Fig. 4), is not needed for binding in telomeric promoters (C as in Fig. 2A).

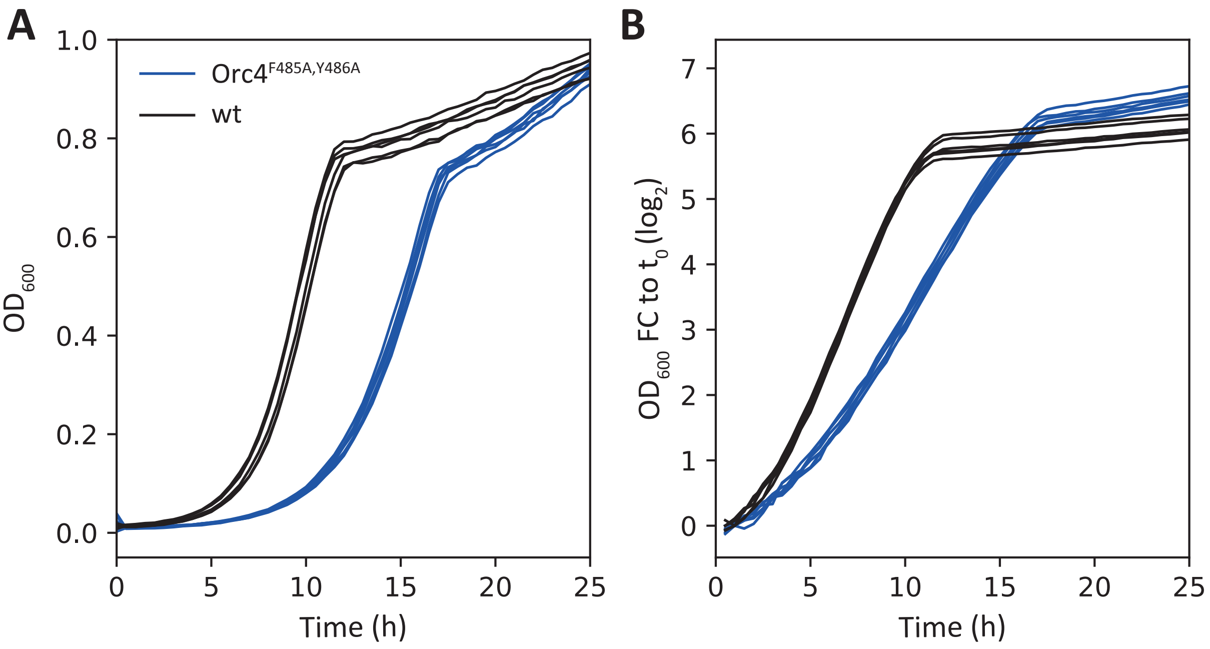


**Figure S3.** **Orc4-IH mutant growth rate deficiency.**
Shown are OD measurements (A) and log2 of the fold change OD measurements (B) for six repeats of the Orc4-IH mutant (F485A,Y486A) and wild type cells. Cells were grown in YPD and constant shaking in 30$^{o}$C.

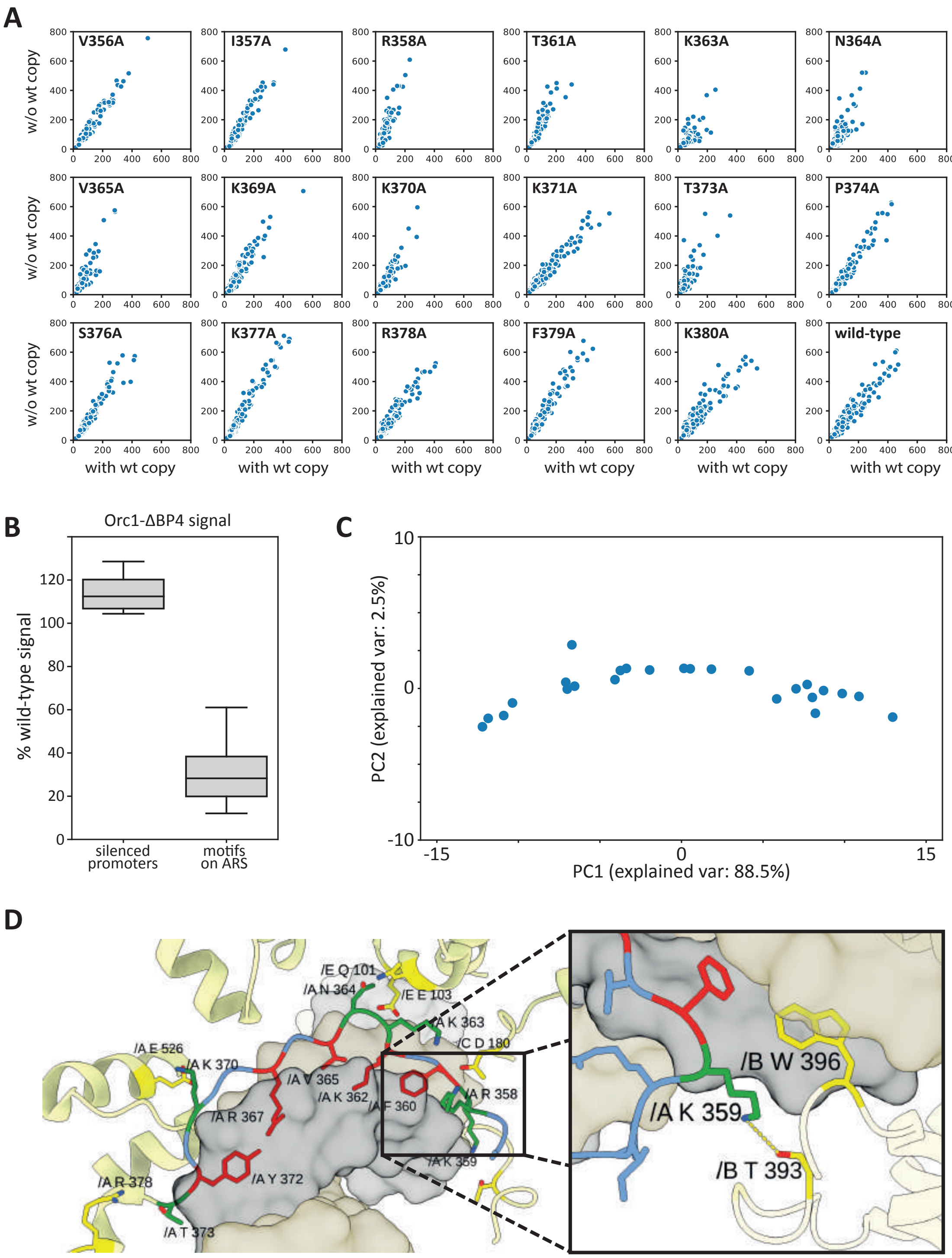


### Figure S4.

A. For each non essential point mutation in Orc1-BP4 and its surrounding residues, two strains were generated: one with an additional Orc1 wild-type copy while the other without. MNase was fused to the mutated Orc1 in both cases, showing a signal only for the mutated copy. Plotted is the binding on ARS consensus sequences in both strains.

B. Binding of Orc1-BP4 deletion in silencing loci vs. ARS motifs.
C. PCA analysis of the binding signal of the Alanine scan mutants on ARS consensus sequences. Plotted are the first and second principal components (PC). The explained variation of each PC is written in the axis titles.
D. Based on the crystal structure, the DNA interacting residues can be classified into two groups: residues that directly interact with DNA as they face the minor groove are marked in red (F360, K362, V365, R367, Y372), and residues that might bind to other parts of the complex are marked in green (R358, K363, N364, K370, T373). For example, zoomed-in view of the K359 residue, which was lethal when mutated to Alanine, interacts with a nearby Threonine from Orc2 (chain B). Accordingly, the K359A effect on binding might be via a network of interactions that is not represented in our simulations as they include only 18 residues and 20 DNA bases.

**Table S1.** Yeast strains used in this study.

| | Strain | Genotype | Background | Parent |
|---|---|---|---|---|
| 1 | ORC1-MNase | MATa his3Δ1 leu2Δ0 met15Δ0 ura3Δ0 **ORC1-N'-3xFlag-MNase** | BY4741[2] | N-SWAT strain from Yofe et al[3]. |
| 2 | ORC2-MNase | MATa his3Δ1 leu2Δ0 met15Δ0 ura3Δ0 **ORC2-N'-3xFlag-MNase** | BY4741[2] | N-SWAT strain from Yofe et al[3]. |
| 3 | ORC4-MNase | MATa his3Δ1 leu2Δ0 met15Δ0 ura3Δ0 **ORC4-N'-3xFlag-MNase** | BY4741[2] | N-SWAT strain from Yofe et al[3]. |
| 4 | ORC1-copy | MATa his3Δ1 leu2Δ0 met15Δ0 ura3Δ0 **bar1Δ** | BY4741[2] | BY4741[2] |
| 5 | Δ bar1, ORC1-copy | MATa his3Δ1 leu2Δ0 met15Δ0 ura3Δ0 **bar1::ORC1-syn** | BY4741[2] | BY4741[2] |
| 6 | ORC1-MNase, Δ bar1 | MATa his3Δ1 leu2Δ0 met15Δ0 ura3Δ0 **bar1Δ ORC1-N'-3xFlag-MNase** | BY4741[2] | Strain 4 |
| 7 | ORC1-MNase, Δ bar1, ORC1-copy | MATa his3Δ1 leu2Δ0 met15Δ0 ura3Δ0 **bar1::ORC1-synCopy ORC1-N'-3xFlag-MNase** | BY4741[2] | BY4741[2] |
| 8 | ORC1-Δ1-188-MNase, Δ bar1 | MATa his3Δ1 leu2Δ0 met15Δ0 ura3Δ0 **bar1Δ ORC1-N'-3xFlag-MNase-Truncation-188** | BY4741[2] | Strain 4 |
| 9 | ORC1-Δ1-290-MNase, Δ bar1 | MATa his3Δ1 leu2Δ0 met15Δ0 ura3Δ0 **bar1Δ ORC1-N'-3xFlag-MNase-Truncation-290** | BY4741[2] | Strain 4 |
| 10 | ORC1-Δ1-320-MNase, Δ bar1 | MATa his3Δ1 leu2Δ0 met15Δ0 ura3Δ0 **bar1Δ ORC1-N'-3xFlag-MNase-Truncation-320** | BY4741[2] | Strain 4 |
| 11 | ORC1-Δ1-340-MNase, Δ bar1 | MATa his3Δ1 leu2Δ0 met15Δ0 ura3Δ0 **bar1Δ ORC1-N'-3xFlag-MNase-Truncation-340** | BY4741[2] | Strain 4 |
| 12 | ORC1-Δ1-188-MNase, Δ bar1, ORC1-copy | MATa his3Δ1 leu2Δ0 met15Δ0 ura3Δ0 **bar1::ORC1-synCopy ORC1-N'-3xFlag-MNase-Truncation-188** | BY4741[2] | Strain 5 |
| 13 | ORC1-Δ1-290-MNase, Δ bar1, ORC1-copy | MATa his3Δ1 leu2Δ0 met15Δ0 ura3Δ0 **bar1::ORC1-synCopy ORC1-N'-3xFlag-MNase-Truncation-290** | BY4741[2] | Strain 5 |
| 14 | ORC1-Δ1-320-MNase, Δ bar1, ORC1-copy | MATa his3Δ1 leu2Δ0 met15Δ0 ura3Δ0 **bar1::ORC1-synCopy ORC1-N'-3xFlag-MNase-Truncation-320** | BY4741[2] | Strain 5 |
| 15 | ORC1-Δ1-340-MNase, Δ bar1, ORC1-copy | MATa his3Δ1 leu2Δ0 met15Δ0 ura3Δ0 **bar1::ORC1-synCopy ORC1-N'-3xFlag-MNase-Truncation-340** | BY4741[2] | Strain 5 |

| | | | | |
|---|---|---|---|---|
| 16 | ORC1-1-375-MNase, Δ bar1, ORC1-Δ1-188-copy | MATa his3Δ1 leu2Δ0 met15Δ0 ura3Δ0 **ORC1-Truncation-188 bar1:: 3xFlag-MNase-ORC1-AA1-375** | BY4741[2] | BY4741[2] |
| 17 | ORC1-188-375-MNase, Δ bar1, ORC1-copy | MATa his3Δ1 leu2Δ0 met15Δ0 ura3Δ0 **bar1:: 3xFlag-MNase-ORC1-AA188-375** | BY4741[2] | BY4741[2] |
| 18 | ORC1-1-357-MNase, Δ bar1, ORC1-copy | MATa his3Δ1 leu2Δ0 met15Δ0 ura3Δ0 **bar1:: 3xFlag-MNase-ORC1-AA1-357** | BY4741[2] | BY4741[2] |
| 19 | ORC1-Δ340-375-MNase, Δ bar1, ORC1-copy | MATa his3Δ1 leu2Δ0 met15Δ0 ura3Δ0 **bar1::ORC1-synCopy ORC1-N'-3xFlag-MNase-Del-340-375** | BY4741[2] | Strain 7 |
| 20 | ORC1-Δ357-375-MNase, Δ bar1, ORC1-copy | MATa his3Δ1 leu2Δ0 met15Δ0 ura3Δ0 **bar1::ORC1-synCopy ORC1-N'-3xFlag-MNase-Del-357-375** | BY4741[2] | Strain 7 |
| 21 | ORC1-MNase, ORC4-F485A-Y486A, Δ bar1 | MATa his3Δ1 leu2Δ0 met15Δ0 ura3Δ0 **bar1Δ ORC1_N'-3xFlag-MNase ORC4-F485A-Y486A** | BY4741[2] | Strain 6 |
| 22 | ORC1-V356A-MNase, Δ bar1 | MATa his3Δ1 leu2Δ0 met15Δ0 ura3Δ0 **bar1Δ ORC1_N'-3xFlag-MNase-V356A** | BY4741[2] | Strain 6 |
| 23 | ORC1-V356A-MNase, Δ bar1, ORC1-copy | MATa his3Δ1 leu2Δ0 met15Δ0 ura3Δ0 **bar1::ORC1-synCopy ORC1_N'-3xFlag-MNase-V356A** | BY4741[2] | Strain 7 |
| 24 | ORC1-I357A-MNase, Δ bar1 | MATa his3Δ1 leu2Δ0 met15Δ0 ura3Δ0 **bar1Δ ORC1_N'-3xFlag-MNase-I357A** | BY4741[2] | Strain 6 |
| 25 | ORC1-I357A-MNase, Δ bar1, ORC1-copy | MATa his3Δ1 leu2Δ0 met15Δ0 ura3Δ0 **bar1::ORC1-synCopy ORC1_N'-3xFlag-MNase-I357A** | BY4741[2] | Strain 7 |
| 26 | ORC1-R358A-MNase, Δ bar1 | MATa his3Δ1 leu2Δ0 met15Δ0 ura3Δ0 **bar1Δ ORC1_N'-3xFlag-MNase-R358A** | BY4741[2] | Strain 7 |
| 27 | ORC1-*R358A*-MNase, Δ bar1, ORC1-copy | MATa his3Δ1 leu2Δ0 met15Δ0 ura3Δ0 **bar1::ORC1-synCopy ORC1_N'-3xFlag-MNase-R358A** | BY4741[2] | Strain 7 |
| 28 | ORC1-*K359A*-MNase, Δ bar1, ORC1-copy | MATa his3Δ1 leu2Δ0 met15Δ0 ura3Δ0 **bar1::ORC1-synCopy ORC1_N'-3xFlag-MNase-K359A** | BY4741[2] | Strain 7 |
| 29 | ORC1-*F360A*-MNase, Δ bar1, ORC1-copy | MATa his3Δ1 leu2Δ0 met15Δ0 ura3Δ0 **bar1::ORC1-synCopy ORC1_N'-3xFlag-MNase-F360A** | BY4741[2] | Strain 7 |
| 30 | ORC1-T361A-MNase, Δ bar1 | MATa his3Δ1 leu2Δ0 met15Δ0 ura3Δ0 **bar1Δ ORC1_N'-3xFlag-MNase-T361A** | BY4741[2] | Strain 6 |
| 31 | ORC1-*T361A*-MNase, Δ bar1, ORC1-copy | MATa his3Δ1 leu2Δ0 met15Δ0 ura3Δ0 **bar1::ORC1-synCopy ORC1_N'-3xFlag-MNase-T361A** | BY4741[2] | Strain 7 |
| 32 | ORC1-*K362A*-MNase, Δ bar1, ORC1-copy | MATa his3Δ1 leu2Δ0 met15Δ0 ura3Δ0 **bar1::ORC1-synCopy ORC1_N'-3xFlag-MNase-K362A** | BY4741[2] | Strain 7 |
| 33 | ORC1-K363A-MNase, Δ bar1 | MATa his3Δ1 leu2Δ0 met15Δ0 ura3Δ0 **bar1Δ ORC1_N'-3xFlag-MNase-K363A** | BY4741[2] | Strain 6 |

| | | | | |
|---|---|---|---|---|
| 34 | ORC1-*K363A*-MNase, Δ bar1, ORC1-copy | MATa his3Δ1 leu2Δ0 met15Δ0 ura3Δ0 **bar1::ORC1-synCopy ORC1_N'-3xFlag-MNase-K363A** | BY4741[2] | Strain 7 |
| 35 | ORC1-N364A-MNase, Δ bar1 | MATa his3Δ1 leu2Δ0 met15Δ0 ura3Δ0 **bar1Δ ORC1_N'-3xFlag-MNase-N364A** | BY4741[2] | Strain 6 |
| 36 | ORC1-*N364A*-MNase, Δ bar1, ORC1-copy | MATa his3Δ1 leu2Δ0 met15Δ0 ura3Δ0 **bar1::ORC1-synCopy ORC1_N'-3xFlag-MNase-N364A** | BY4741[2] | Strain 7 |
| 37 | ORC1-V365A-MNase, Δ bar1 | MATa his3Δ1 leu2Δ0 met15Δ0 ura3Δ0 **bar1Δ ORC1_N'-3xFlag-MNase-V365A** | BY4741[2] | Strain 6 |
| 38 | ORC1-*V365A*-MNase, Δ bar1, ORC1-copy | MATa his3Δ1 leu2Δ0 met15Δ0 ura3Δ0 **bar1::ORC1-synCopy ORC1_N'-3xFlag-MNase-V365A** | BY4741[2] | Strain 7 |
| 39 | ORC1-*R367A*-MNase, Δ bar1, ORC1-copy | MATa his3Δ1 leu2Δ0 met15Δ0 ura3Δ0 **bar1::ORC1-synCopy ORC1_N'-3xFlag-MNase-R367A** | BY4741[2] | Strain 7 |
| 40 | ORC1-K369A-MNase, Δ bar1 | MATa his3Δ1 leu2Δ0 met15Δ0 ura3Δ0 **bar1Δ ORC1_N'-3xFlag-MNase-K369A** | BY4741[2] | Strain 6 |
| 41 | ORC1-*K369A*-MNase, Δ bar1, ORC1-copy | MATa his3Δ1 leu2Δ0 met15Δ0 ura3Δ0 **bar1::ORC1-synCopy ORC1_N'-3xFlag-MNase-K369A** | BY4741[2] | Strain 7 |
| 42 | ORC1-K370A-MNase, Δ bar1 | MATa his3Δ1 leu2Δ0 met15Δ0 ura3Δ0 **bar1Δ ORC1_N'-3xFlag-MNase-K370A** | BY4741[2] | Strain 6 |
| 43 | ORC1-*K370A*-MNase, Δ bar1, ORC1-copy | MATa his3Δ1 leu2Δ0 met15Δ0 ura3Δ0 **bar1::ORC1-synCopy ORC1_N'-3xFlag-MNase-K370A** | BY4741[2] | Strain 7 |
| 44 | ORC1-K371A-MNase, Δ bar1 | MATa his3Δ1 leu2Δ0 met15Δ0 ura3Δ0 **bar1Δ ORC1_N'-3xFlag-MNase-K371A** | BY4741[2] | Strain 6 |
| 45 | ORC1-*K371A*-MNase, Δ bar1, ORC1-copy | MATa his3Δ1 leu2Δ0 met15Δ0 ura3Δ0 **bar1::ORC1-synCopy ORC1_N'-3xFlag-MNase-K371A** | BY4741[2] | Strain 7 |
| 46 | ORC1-*Y372A*-MNase, Δ bar1, ORC1-copy | MATa his3Δ1 leu2Δ0 met15Δ0 ura3Δ0 **bar1::ORC1-synCopy ORC1_N'-3xFlag-MNase-Y372A** | BY4741[2] | Strain 7 |
| 47 | ORC1-T373A-MNase, Δ bar1 | MATa his3Δ1 leu2Δ0 met15Δ0 ura3Δ0 **bar1Δ ORC1_N'-3xFlag-MNase-T373A** | BY4741[2] | Strain 6 |
| 48 | ORC1-*T373A*-MNase, Δ bar1, ORC1-copy | MATa his3Δ1 leu2Δ0 met15Δ0 ura3Δ0 **bar1::ORC1-synCopy ORC1_N'-3xFlag-MNase-T373A** | BY4741[2] | Strain 7 |
| 49 | ORC1-P374A-MNase, Δ bar1 | MATa his3Δ1 leu2Δ0 met15Δ0 ura3Δ0 **bar1Δ ORC1_N'-3xFlag-MNase-P374A** | BY4741[2] | Strain 6 |
| 50 | ORC1-*P374A*-MNase, Δ bar1, ORC1-copy | MATa his3Δ1 leu2Δ0 met15Δ0 ura3Δ0 **bar1::ORC1-synCopy ORC1_N'-3xFlag-MNase-P374A** | BY4741[2] | Strain 7 |
| 51 | ORC1-F375A-MNase, Δ bar1 | MATa his3Δ1 leu2Δ0 met15Δ0 ura3Δ0 **bar1Δ ORC1_N'-3xFlag-MNase-F375A** | BY4741[2] | Strain 6 |
| 52 | ORC1-S376A-MNase, Δ bar1 | MATa his3Δ1 leu2Δ0 met15Δ0 ura3Δ0 **bar1Δ ORC1_N'-3xFlag-MNase-S376A** | BY4741[2] | Strain 6 |

| | | | | |
|---|---|---|---|---|
| 53 | ORC1-*S376A*-MNase, Δ bar1, ORC1-copy | MATa his3Δ1 leu2Δ0 met15Δ0 ura3Δ0 **bar1::ORC1-synCopy ORC1_N'-3xFlag-MNase-S376A** | BY4741[2] | Strain 7 |
| 54 | ORC1-K377A-MNase, Δ bar1 | MATa his3Δ1 leu2Δ0 met15Δ0 ura3Δ0 **bar1Δ ORC1_N'-3xFlag-MNase-K377A** | BY4741[2] | Strain 6 |
| 55 | ORC1-*K377A*-MNase, Δ bar1, ORC1-copy | MATa his3Δ1 leu2Δ0 met15Δ0 ura3Δ0 **bar1::ORC1-synCopy ORC1_N'-3xFlag-MNase-K377A** | BY4741[2] | Strain 7 |
| 56 | ORC1-R378A-MNase, Δ bar1 | MATa his3Δ1 leu2Δ0 met15Δ0 ura3Δ0 **bar1Δ ORC1_N'-3xFlag-MNase-R378A** | BY4741[2] | Strain 6 |
| 57 | ORC1-*R378A*-MNase, Δ bar1, ORC1-copy | MATa his3Δ1 leu2Δ0 met15Δ0 ura3Δ0 **bar1::ORC1-synCopy ORC1_N'-3xFlag-MNase-R378A** | BY4741[2] | Strain 7 |
| 58 | ORC1-F379A-MNase, Δ bar1 | MATa his3Δ1 leu2Δ0 met15Δ0 ura3Δ0 **bar1Δ ORC1_N'-3xFlag-MNase-F379A** | BY4741[2] | Strain 6 |
| 59 | ORC1-*F379A*-MNase, Δ bar1, ORC1-copy | MATa his3Δ1 leu2Δ0 met15Δ0 ura3Δ0 **bar1::ORC1-synCopy ORC1_N'-3xFlag-MNase-F379A** | BY4741[2] | Strain 7 |
| 60 | ORC1-K380A-MNase, Δ bar1 | MATa his3Δ1 leu2Δ0 met15Δ0 ura3Δ0 **bar1Δ ORC1_N'-3xFlag-MNase-K380A** | BY4741[2] | Strain 6 |
| 61 | ORC1-*K380A*-MNase, Δ bar1, ORC1-copy | MATa his3Δ1 leu2Δ0 met15Δ0 ura3Δ0 **bar1::ORC1-synCopy ORC1_N'-3xFlag-MNase-K380A** | BY4741[2] | Strain 7 |
| 62 | ORC1-Δ320-375-MNase, Δ bar1, ORC1-copy | MATa his3Δ1 leu2Δ0 met15Δ0 ura3Δ0 **bar1::ORC1-synCopy ORC1-N'-3xFlag-MNase-Del-320-375** | BY4741[2] | Strain 7 |
| 63 | ORC1-Δ320-340-MNase, Δ bar1, ORC1-copy | MATa his3Δ1 leu2Δ0 met15Δ0 ura3Δ0 **bar1::ORC1-synCopy ORC1-N'-3xFlag-MNase-Del-320-340** | BY4741[2] | Strain 7 |
| 64 | ORC1-Δ320-340-MNase, Δ bar1 | MATa his3Δ1 leu2Δ0 met15Δ0 ura3Δ0 **bar1Δ ORC1-N'-3xFlag-MNase-Del-320-340** | BY4741[2] | Strain 6 |
| 65 | ORC1-Δ320-340-MNase, ORC4-F485A-Y486A, ORC1-copy, Δ bar1 | MATa his3Δ1 leu2Δ0 met15Δ0 ura3Δ0 **bar1::ORC1-synCopy ORC1-N'-3xFlag-MNase-Del-320-340 ORC4-F485A-Y486A** | BY4741[2] | Strain 63 |

**Data S1.** ORC1- synonymous copy sequence (with ORC1 5’UTR and 3’UTR):

AAGAAAACTGCATAGGCGGCAAATTCAGCCTAAAAGTTTCCAGAAGCAGGAACTCATTCCCTATTGATT
AATACTCATTACAAAAACCACAATAGAGTAGATAAG**ATGGCCAAAACTTTAAAGGATTTGCAAGGATGG**
**GAAATTATTACCACAGATGAGCAAGGAAATATTATAGATGGAGGCCAGAAAAGACTAAGAAGAAGAGGT**
**GCCAAAACGGAACATTACTTGAAGAGGAGTTCGGACGGCATTAAATTAGGTCGAGGGGATAGTGTAGTT**
**ATGCATAATGAAGCAGCTGGTACTTATTCAGTATACATGATTCAAGAGCTGCGTCTTAACACATTGAAC**
**AACGTGGTGGAACTGTGGGCACTAACTTATTTAAGATGGTTTGAAGTTAATCCACTGGCGCACTACAGA**
**CAATTTAACCCAGATGCAAATATCCTAAATAGACCTTTGAACTATTATAATAAGTTGTTTTCAGAAACA**
**GCAAATAAAAACGAATTGTATTTGACGGCTGAATTGGCGGAATTACAATTATTTAATTTTATTAGAGTA**
**GCTAATGTTATGGATGGTTCTAAGTGGGAGGTCTTAAAGGGAAATGTAGATCCGGAAAGAGATTTTACT**
**GTTAGATACATTTGTGAACCGACAGGCGAGAAGTTTGTAGATATTAACATAfGAAGACGTTAAAGCCTA**
**TATAAAGAAGGTAGAACCCCGCGAAGCTCAAGAGTATTTAAAAGATCTCACGTTfACCATCAAAGAAAA**
**AAGAAATTAAAAGGGGACCTCAGAAGAAAGACAAAGCAACGCAGACTGCTCAAATATCTGATGCAGAGA**
**CTAGGGCTACGGACATTACGGACAATGAGGACGGAAATGAAGATGAATCGTCAGATTATGAATCCCCAT**
**CTGATATTGATGTCTCAGAAGATATGGATAGTGGTGAAATATCAGCCGACGAGTTAGAAGAAGAAGAGG**
**ATGAAGAGGAAGACGAAGATGAAGAGGAAAAAGAGGCTAGACACACAAATTCACCTAGAAAGAGGGGAC**
**GAAAAATAAAGCTTGGAAAAGATGACATAGACGCTAGTGTCCAACCTCCTCCCAAAAAGAGAGGGAGGA**
**AACCAAAAGACCCTTCTAAACCTCGTCAGATGCTACTAATATCAAGTTGTAGGGCTAATAATACACCTG**
**TAATCAGAAAATTTACTAAGAAAAATGTTGCGCGTGCCAAAAAAAAATATACTCCGTTTAGCAAAAGAT**
**TCAAGTCAATTGCCGCTATTCCTGATTTGACATCCTTGCCAGAATTCTATGGGAACTCTTCCGAATTGA**
**TGGCATCTCGTTTCGAAAATAAGTTAAAAACTACCCAAAAACACCAGATAGTAGAAACTATTTTTTCGA**
**AAGTGAAAAAGCAGCTTAATAGTTCTTATGTGAAGGAAGAGATTTTAAAGAGTGCCAATTTTCAAGATT**
**ACTTACCTGCTAGGGAAAATGAATTTGCCTCAATTTACTTAAGCGCTTACTCTGCTATAGAATCAGATT**
**CAGCTACTACCATATATGTAGCTGGAACACCCGGTGTTGGCAAAACATTGACGGTTAGAGAAGTAGTCA**
**AAGAGTTACTTAGTTCATCAGCTCAGCGTGAAATACCAGATTTCCTATACGTAGAAATTAATGGACTAA**
**AAATGGTCAAGCCTACTGATTGCTATGAAACACTTTGGAACAAAGTCTCCGGGGAAAGGTTGACCTGGG**
**CTGCTTCGATGGAAAGCTTAGAGTTCTATTTCAAAAGAGTCCCAAAAAACAAAAAAAAGACAATTGTTG**
**TTTTATTAGACGAACTGGACGCCATGGTTACTAAATCTCAGGATATAATGTACAATTTCTTTAATTGGA**
**CAACATATGAAAATGCAAAACTCATTGTGATAGCAGTAGCAAATACGATGGATCTACCGGAGCGTCAAT**
**TAGGGAATAAGATTACTAGTCGCATCGGATTTACTAGGATTATGTTTACTGGGTATACACATGAAGAAT**
**TGAAAAATATTATTGACTTAAGACTAAAGGGTTTGAATGATTCTTTTTTTTATGTGGATACCAAAACGG**
**GAAATGCCATTCTAATAGACGCAGCAGGTAATGATACAACTGTTAAACAAACATTACCGGAAGACGTCA**
**GAAAGGTCAGATTGAGAATGTCTGCTGACGCCATCGAGATAGCTTCGAGGAAGGTCGCATCTGTATCAG**
**GTGATGCACGTCGAGCACTCAAAGTTTGTAAAAGAGCCGCTGAAATTGCCGAGAAACACTATATGGCAA**
**AACACGGTTACGGTTACGACGGCAAAACCGTTATTGAAGACGAAAATGAAGAGCAGATTTATGATGATG**
**AAGATAAAGATTTGATAGAATCAAATAAGGCAAAGGACGATAATGACGACGATGATGATAATGATGGTG**
**TACAGACGGTCCATATTACTCATGTCATGAAAGCACTGAACGAGACACTTAATTCTCATGTGATTACTT**
**TTATGACAAGATTGAGTTTTACGGCCAAACTCTTTATATATGCATTGTTAAATTTGATGAAAAAAAACG**
**GATCTCAGGAGCAAGAATTAGGTGATATCGTTGACGAAATAAAATTACTCATCGAAGTCAATGGGAGTA**
**ACAAATTTGTAATGGAAATAGCCAAGACATTATTTCAACAAGGATCAGATAATATTTCTGAGCAACTTA**
**GGATCATTTCTTGGGATTTTGTATTAAATCAGCTTTTAGACGCAGGTATACTTTTTAAACAGACTATGA**
**AAAATGATCGTATCTGTTGTGTAAAATTGAATATTAGTGTTGAAGAGGCAAAAAGGGCTATGAATGAGG**
**ATGAAACTTTACGAAACTTGTAG**ATTCGGTTTTTATTATTCATGACCTAGCATACACATACATATACCT
ACATAGTAGCGCATTTATCCAAAACATACGATATTGTGGATGTACATACCTTCTATATCTCCTTAAAGC
TATTGTGTAGaCTTGATTTAAAATATGCTAACGCCA
